# Rudhira/BCAS3 couples microtubules and intermediate filaments to promote cell migration for angiogenic remodeling

**DOI:** 10.1101/381384

**Authors:** Divyesh Joshi, Maneesha S. Inamdar

## Abstract

Blood vessel formation requires endothelial cell (EC) migration that depends on dynamic remodeling of the cytoskeleton. Rudhira/Breast Carcinoma Amplified Sequence 3 (BCAS3) is a cytoskeletal protein essential for EC migration and sprouting angiogenesis during mouse development and implicated in metastatic disease. Here, we report that Rudhira mediates cytoskeleton organization and dynamics during EC migration. Rudhira binds to both microtubules and Vimentin intermediate filaments (IFs) and stabilizes microtubules. Rudhira depletion impairs cytoskeletal crosstalk, microtubule stability and hence focal adhesion disassembly. The BCAS3 domain of Rudhira is necessary and sufficient for microtubule-IF crosslinking and cell migration. Pharmacologically restoring microtubule stability rescues gross cytoskeleton organization and angiogenic sprouting in Rudhira depleted cells. Our study identifies the novel and essential role of Rudhira in cytoskeletal crosstalk and assigns function to the conserved BCAS3 domain. Targeting Rudhira could allow tissue-restricted cytoskeleton modulation to control cell migration and angiogenesis in development and disease.

## Introduction

Cell migration in physiological or pathological contexts depends on co-ordinated changes in the cytoskeleton and cell-matrix adhesions. Directed endothelial cell (EC) migration is an important pre-requisite for developmental as well as pathological angiogenesis. ECs respond to molecular or mechanical cues in the dynamically changing microenvironment as they move to target tissues for sprouting and angiogenic remodeling. While the fundamental cytoskeletal machinery operates in ECs, few EC-specific cytoskeletal modulators are known. Perturbing the cytoskeleton results in dramatic loss of EC function. For example, non-centrosomal microtubules (MTs) and Vimentin IFs have recently been shown to have an indispensable role in sprouting angiogenesis [1, 2]. Further, disruption of either plus or minus ends of MTs, can inhibit MT-actin crosstalk, adhesion dynamics and thereby EC sprouting [3]. Regulation of cytoskeletal interactions is likely to be important in developmental as well as tumour angiogenesis.

While MTs and IFs can interact directly, several molecules are known to bridge cytoskeletal components. Cytolinkers of the plakin family are well characterized and connect MTs, IFs, actin filaments and plasma membrane components. The importance of cytoskeletal crosstalk is evident from the early postnatal lethality of mice lacking the prototype cytolinker Plectin [4]. Many ubiquitously expressed molecules like the MT motor Kinesin and tumor suppressor APC are critical for MT-IF crosslinking in fibroblasts and migrating astrocytes respectively [5, 6]. Recent elegant studies show that while MTs are essential for Vimentin IF assembly, Vimentin IFs provide memory for MT cytoskeleton regrowth highlighting the significance of the crosstalk and the positive feedback interaction between these two cytoskeletal components [7, 8].

Cytoskeletal components play a critical role in establishing cell polarity, force generation and regulation of adhesion complex components during cell migration. Close and intricate interactions between actin filaments, microtubules (MTs), intermediate filaments (IFs) and cytoskeleton-associated proteins bring about dynamic reorganization of cell shape and focal adhesions (FAs), essential for directed cell migration. Association with Vimentin IFs stabilizes MTs against depolymerising stresses and shrinkage, likely owing to the ten-fold slower turnover rate of Vimentin IFs [7]. In addition, MTs have been proposed to grow along Vimentin IFs, although the bridging components in this process remain elusive [7]. The physiological significance of this interaction is also unclear, in part, owing to the full-term survival of *vimentin* knockout mouse and the likely redundancy in IF functions [9]. Several cytoskeleton-associated molecules like the MT plus-end tracking proteins (+TIPs) EB1 and CLIP170 and spectraplackin family member ACF7 cross-bridge MTs and actin and guide MT growth to FAs for FA turnover and persistent migration [10].

Depletion of individual cytoskeleton components, associated proteins or cytolinkers often disrupts overall cytoskeleton architecture and dynamics, perturbing cell migration and adhesion. Multiple and/or redundant roles of the molecules involved as well as context-dependent responses make it challenging to decipher global and tissue-specific mechanisms that regulate this process. Identifying additional molecular components could help unravel mechanisms to control or promote cell migration in desired contexts.

Rudhira/BCAS3 (Breast Carcinoma Amplified Sequence 3) is a cytoskeletal protein essential for mouse developmental angiogenesis and implicated in tumor metastasis [11–13]. Rudhira binds to MTs and intermediate filaments (IFs) and promotes directional cell migration [11]. In this study we examine the mechanism by which Rudhira controls cytoskeletal remodeling during cell migration. We show that Rudhira directly associates with MTs and IFs for MT-IF crosstalk, MT stability and dynamics and thereby FA turnover and cell migration, through its conserved BCAS3 domain. Our study provides new insights into the mechanism of cytoskeletal crosslinking and reorganization during cell migration, which will help understand physiological and pathological angiogenesis.

## Results

### Rudhira is required for gross cytoskeletal organization

We reported earlier that Rudhira/BCAS3 interacts with microtubules (MT) and intermediate filaments (IFs) and is required for actin reorganization for directional endothelial cell (EC) migration. Rudhira depletion deregulates several cellular and molecular processes critical for sprouting angiogenesis *in vitro* and *in vivo*, including cell adhesion and invasion [12]. To explore the possible mechanisms by which Rudhira functions in cell migration, we examined the effect of Rudhira depletion (knockdown, KD) on cytoskeletal organization as compared to the non-silencing control (NS) in mouse Saphenous Vein Endothelial Cell line (SVEC). Immunolocalization showed that unlike in control, in KD cells MTs were not aligned towards and appeared bent at the cell periphery while Vimentin IFs were fewer and not extended but present only in the perinuclear region (Figure 1A). KD cells also had thick actin bundles at the cell cortex (Figure 1B) in addition to increased stress fibres, suggesting aberrant cell-substratum adhesion [14]. However, protein levels of the cytoskeletal components were unaltered (Figure 1C).

**Figure 1.**
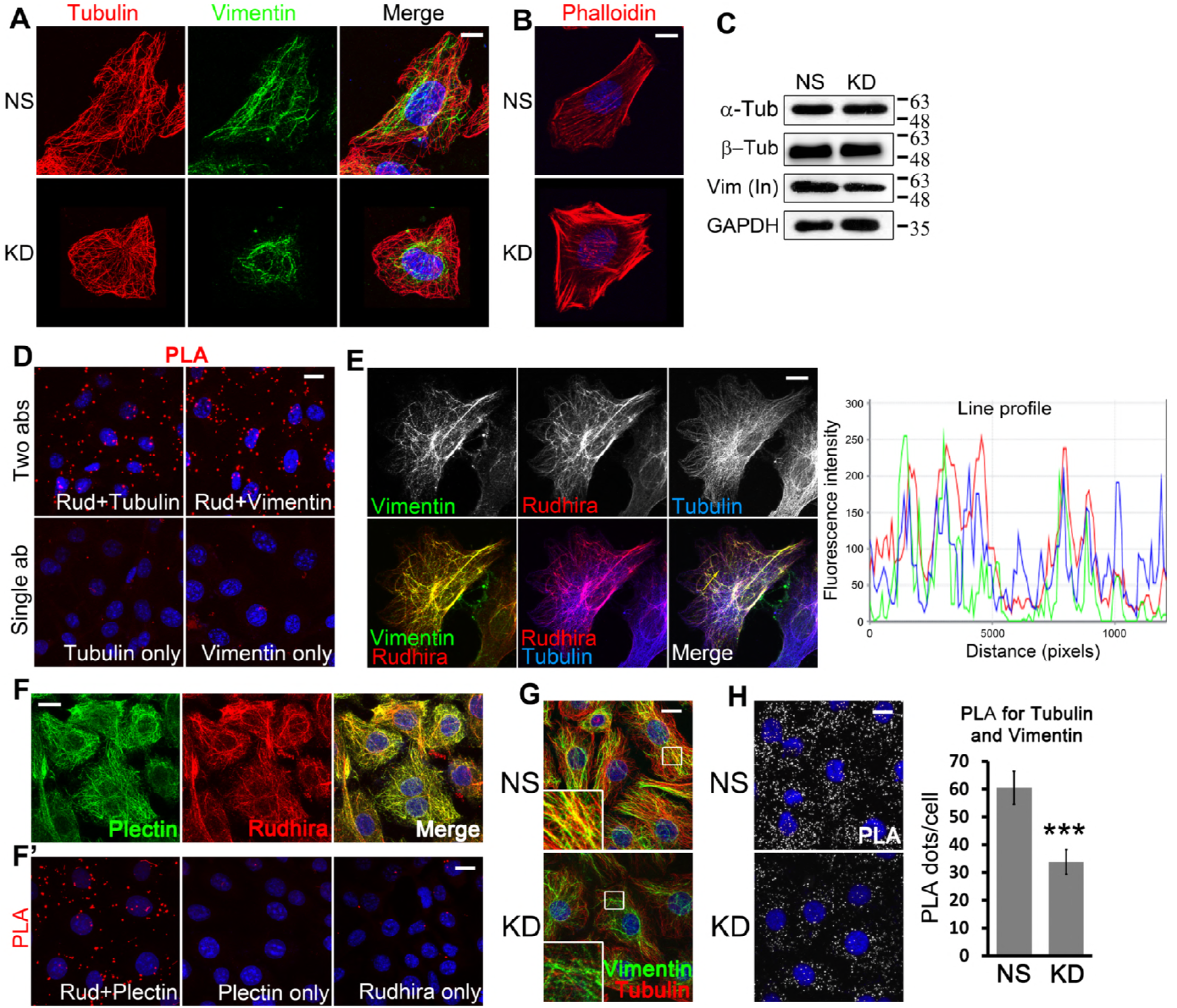
Rudhira interacts with and controls crosstalk between microtubules and intermediate filaments. (A, B) NS and KD cells were co-stained for cytoskeleton markers, Tubulin and Vimentin (A) or Phalloidin (B) to detect gross cytoskeleton organization. (C) Immunoblot to detect the levels of cytoskeletal proteins. (D) Direct interaction between Rudhira and Tubulin or Vimentin in wild type SVECs analysed by Proximity Ligation Assay (PLA). Single antibody stained cells were taken as negative controls. (E) Relative localization of Vimentin IFs, Rudhira and MTs was performed by triple immunostaining in wild type SVECs. Line profile shows the fluorescence intensity peaks for the three colours along the yellow arrow. (F, F’) Relative association of Rudhira with the cytolinker Plectin by immunofluorescence (F) or PLA (F’). Single antibody stained cells were taken as negative controls for PLA. (G) NS or KD cells were analysed for coalignment of Vimentin and MTs by co-immunofluorescence. Boxed regions are magnified in the insets. (H) Vimentin and MT association by Proximity Ligation Assay (PLA). Graph shows the quantitation of PLA dots per cell indicating extent of interaction. Error bars indicate standard error of mean (SEM). Results shown are a representative of at least three independent experiments with at least three biological replicates taken into account. Statistical analysis was carried out using one-way ANOVA. Scale Bar: (A, B, E) 10 μm, (D, F, F’, G, H) 10 μm. *p<0.05, **p<0.01, ***p<0.001.

### Rudhira directly interacts with and bridges IFs and MTs

The intricate association of cytoskeletal components is dynamically regulated during cell migration. MTs and Vimentin IFs are coaligned in mesenchymal cells for efficient migration. While initially Vimentin IFs form along MTs, later these filaments provide a template for MT growth [7]. Further, IFs organize primarily by MT-dependent transport and actin-dependent flow, suggestive of extensive cytoskeletal crosstalk and cross-regulation during cell migration [8]. Rudhira interacts with Tubulin and Vimentin and localizes to MTs and IFs. To test the likely direct interaction of Rudhira with MTs and IFs *in vivo*, we used Proximity Ligation Assay (PLA), which detects interaction at single molecule resolution. Rudhira associated with both Tubulin and Vimentin, suggesting direct interactions of Rudhira with these cytoskeleton components within the cells (Figure 1D). In addition, triple immunofluorescence analysis showed that Rudhira associates with MTs at sites often overlapping with Vimentin IFs suggesting that these interactions may be regulated by local factors or Tubulin or Vimentin properties (Figure 1E and line profile). These data indicate that Rudhira may simultaneously associate with MTs and IFs and IF association of Rudhira may favour its binding to MTs.

In agreement with its proposed role of bridging cytoskeletal components, we observed high overlap of Rudhira with the known cytolinker protein Plectin (Figure 1F). PLA between Rudhira and Plectin confirmed their interaction and suggested that they may have similar function *in vivo* (Figure 1F’). Expectedly, MT-IF association *in vivo* was dramatically reduced in Rudhira depleted cells, as detected by double immunofluorescence (Figure 1G) and confirmed by PLA (Figure 1H). Thus, Rudhira is critical for MT-IF bridging in ECs.

### Rudhira governs the association and dynamics of microtubules and Vimentin intermediate filaments in endothelial cells

The crosstalk between IFs and MTs is essential for efficient EC migration [7, 15]. Hence, we tested for the association of IFs and MTs in live cells in low density cultures, where cells are in a migratory state. Live imaging of MTs using Silico-Rhodamine conjugated Docetaxel (SiR-Tubulin) showed that MTs grew radially towards the cell periphery and were stabilized there in control cells (NS), whereas KD had fewer MTs at the cell periphery and they often started to bend before reaching the periphery (red asterisk in Figure S1A and Video S1). To test MT-IF crosstalk we transiently expressed Vimentin-GFP and incubated cells with SiR-Tubulin. Like the endogenous Vimentin (Figure 1A, G), Vimentin-GFP filaments were less extended in KD cells, resulting in reduced alignment with MTs and perturbed dynamics (Figure 2A and Video S2). These data suggest that Rudhira is required for cytoskeletal crosstalk and organization for cell migration. Rudhira may also have a role in promoting the assembly of or stabilizing IFs.

**Figure 2.**
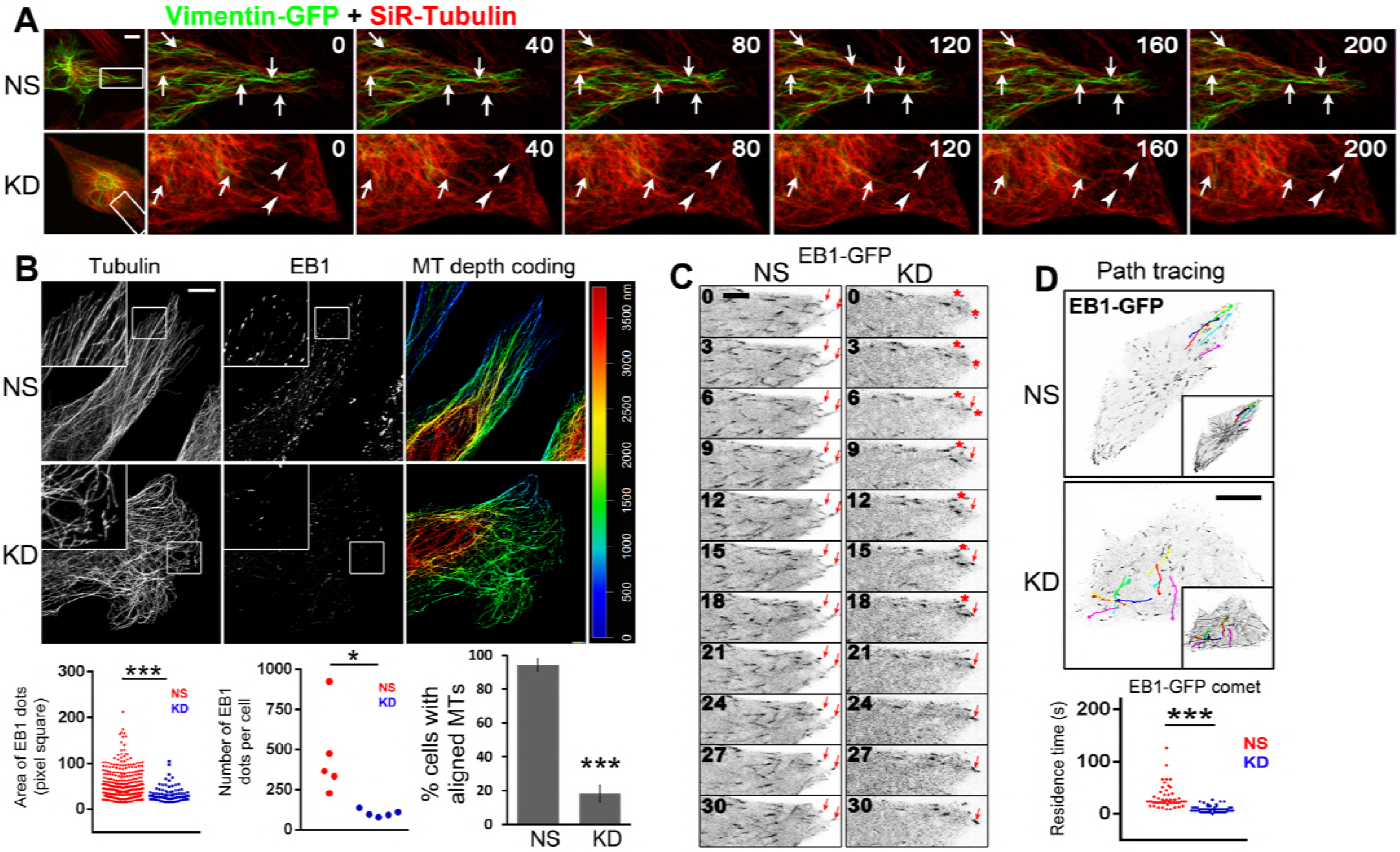
Rudhira is required for MT-Vimentin IF association and dynamics in migrating endothelial cells. (A) Time-lapse images of SVEC NS and KD cells transiently transfected with Vimentin-GFP and stained with SiR-Tubulin, imaged at 10-s intervals. Arrows indicate persistence of coaligned Vimentin IFs and MTs towards the cell periphery while arrowheads indicate the absence of Vimentin IFs and bent MTs before reaching the cell periphery in live migrating cells. (B) Super-resolution imaging after immunostaining NS and KD cells for β-Tubulin and EB1 to detect cell peripheral MT architecture and growing MTs. Panel to the right shows depth coding to detect MTs at cell periphery, the sites near cell-matrix contacts. The graphs show area of EB1 dots (2329 EB1 dots for NS and 514 EB1 dots for KD, over 5 images each), number of EB1 dots per cell (5 images each) and percentage of cells with aligned MTs (40 cells for NS and 46 cells for KD). Boxed regions are magnified in the insets. (C) Time-lapse images of SVEC NS and KD cells transiently transfected with EB1-GFP and imaged at 3-s intervals. Arrows indicate persistence of aligned EB1 positive MT growing end in NS and not in KD while red asterisk indicates a MT end not stabilized at the cell periphery. 20 live cells each of NS and KD were imaged. (D) Overlay of EB1-GFP tracks and their time-projection (insets) in NS and KD. Time-lapse images of a total of 50 randomly selected EB1-GFP comets from 5 cells each were analyzed manually for calculating residence-time at the cell periphery shown in the graph. Error bars indicate standard error of mean (SEM). Results shown are a representative of at least three independent experiments with at least three biological replicates taken into account. Statistical analysis was carried out using one-way ANOVA. Scale Bar: (A, D) 10 μm, (B) 5 μm, (C) 1 μm. *p<0.05, **p<0.01, ***p<0.001.

The MT cytoskeleton is a highly dynamic macromolecular assembly, with a turnover rate of 5-15 min. MTs govern cell polarity and along with actin also control IF organization during migration. Controlled MT dynamics and stability contribute to cell migration by release of cell-ECM contacts and polarized asymmetric distribution of vesicles. Consistent with the earlier observation (Figure 1A), super-resolution microscopy of KD cells showed defective MT arrays with MTs often failing to reach the cell periphery as observed by depth-coding of MTs (Figure 2B). Further, MTs in KD cells seemed to cross over each other, indicating undirected growth, unlike in control cells which displayed aligned MTs near periphery. In addition, KD cells showed reduced co-staining for the +TIP, EB1, indicative of fewer growing MTs (Figure 2B).

To test whether impaired migration of KD cells is due to defects in MT growth, we assessed EB1-GFP transfected control and KD cells by live imaging. Rudhira-depleted cells had fewer EB1-GFP positive MTs (Figure 2C and Video S3). +TIPs bind to and stabilize MTs at FAs for a time period of more than 15 s. The EB1 comets in KD cells appeared to be smaller in size and shorter-lived than those in control cells (Figure 2B, C, D and Video S3). Time-projected (for 60 s) images (see Methods) also showed that MT growth in KD cells followed a criss-cross pattern towards the cell periphery, as compared to the straight linear growth of MTs in control, as judged by EB1-GFP movement in live cells (Figure 2C, D and Video S4). Unlike MTs in controls which grew radially towards and were stabilized at the periphery, MTs in KD cells were rarely stabilized and often started to bend before reaching the cell periphery. (Figure 2D and Video S3, S4). This suggests that MTs in KD cells encounter a physical constraint, likely thick actin stress fibres (Figure 1B), which may prevent their growth to the periphery, the site of cell-matrix adhesions.

### Rudhira associates with and stabilizes microtubules

The differential association and dissociation of Microtubule-associated Proteins (MAPs) and cytoskeletal components modulates MT stability, essential for cell migration. Alpha-tubulin acetylation or detyrosination (Glu) classically marks stable MTs [16]. Acetylation also provides mechanical resistance to breakage [17]. The cytoskeleton in KD cells was grossly disorganized, and our data indicated a role of Rudhira in MT and Vimentin IF crosstalk. Since Vimentin preferentially associates with stable MTs [18] and functions to stabilize MTs, we tested whether binding of Rudhira contributes to MT stability. Stable MTs are oriented towards the leading edge of a migrating cell and due to their higher affinity for MT motors *in vitro* are considered to maintain directional migration by polarized delivery of vesicles [19]. Immunolocalization for acetylated (Ac) tubulin in a scratched EC monolayer 2 h after wounding, showed that compared to control, KD cells had fewer stable MTs that did not reach the leading edge (Figure 3A). Immunoblot also showed significant decrease in Ac- and Glu- α-tubulin levels, indicating that Rudhira depletion destabilized MTs (Figure 3B). This was confirmed by treatment with MT-depolymerising drug and cold treatment. As compared to controls, MTs in Rudhira-depleted cells were more sensitive to both MT depolymerization stresses (Figure 3C, C’). 10 μM Nocodazole caused complete MT depolymerization in both control and KD cells. However low concentrations of Nocodazole (4 nM to 400 nM) depolymerise dynamic but not stable MTs in a dose-dependent manner. KD cells were more sensitive to 10 nM Nocodazole and showed a dramatic reduction in MT number as compared to controls, which showed a well-organized MT-array with little apparent reduction in MT numbers (Figure 3C).

**Figure 3.**
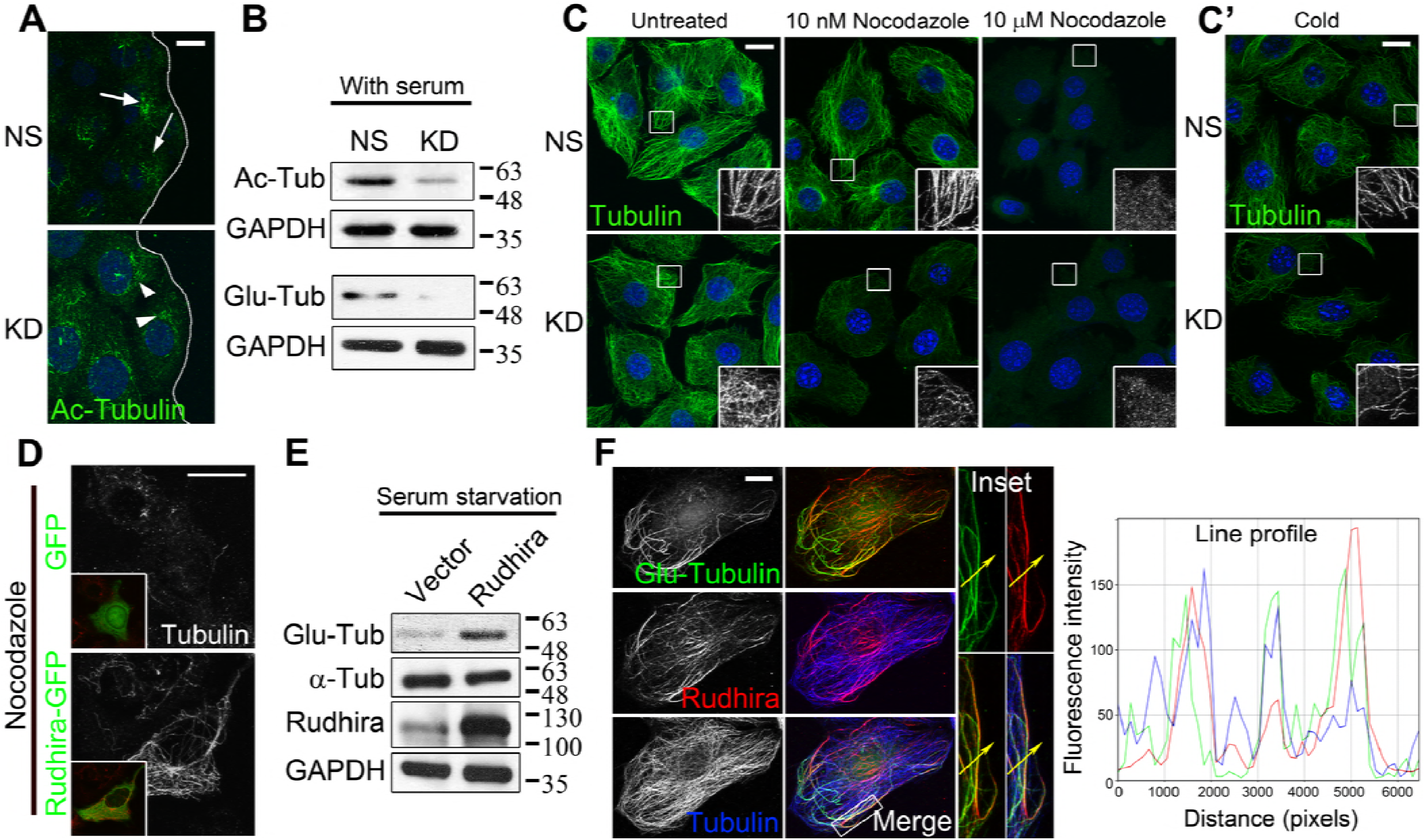
Rudhira is essential for microtubule stability. (A) Localization for acetylated MTs was analyzed in NS and KD cells by immunostaining post a scratch-wound healing assay. Arrows point to acetylated MTs towards the leading edge and arrowheads to acetylated MTs distributed all over the cell. Dotted line represents the wound margin. (B) Acetylated-Tubulin, Glu-Tubulin levels were analysed by immunoblot. (C, C’) NS and KD cells were treated with the indicated dosages of Nocodazole for 30 min (C) or cold PBS at 4 °C (C’) and MTs were analyzed by immunostaining for β-Tubulin. Boxed regions in (c, c’) are magnified in the insets. (D) SVEC cells were transiently transfected with GFP or Rudhira-GFP (inset), treated with Nocodazole and MTs were analyzed by immunostaining for Tubulin. (E) Immunoblot analysis for Glu-Tubulin levels post 48 h serum starvation in HEK293 cells overexpressing Rudhira. (F) Relative localization of detyrosinated MTs (Glu-Tubulin), Rudhira and total MTs (Tubulin) was performed by triple immunostaining in wild type SVECs. Line profile shows the fluorescence intensity peaks for the three colours along the yellow arrow in the inset (magnified boxed region). Error bars indicate standard error of mean (SEM). Results shown are a representative of three independent experiments. Statistical analysis was carried out using one-way ANOVA. Scale Bar: (A, C, C’) 20 μm, (D, F) 10 μm.

Treatment of cells that overexpress Rudhira with MT-depolymerising doses of Nocodazole showed that their MTs are Nocodazole-resistant, as compared to control, where most MTs were depolymerised (Figure 3D, S1B). Further, Glu-tubulin levels were increased (Figure 3E) and the stable MTs were often associated with Rudhira as seen by immunolocalization (Figure S1C). Triple immunofluorescence analysis showed that Rudhira had a preferential association with detyrosinated MTs (Figure 3F and line profile). Thus, like Vimentin IFs, Rudhira binds to and stabilizes MTs and promotes MT-IF association likely leading to MT stability.

### Rudhira-depleted cells have large focal adhesions

MT dynamics and stability have been well studied in the context of cell migration. Cells adhere to the ECM ligands via focal adhesions (FAs) assembled on the cell-peripheral ends of actin stress fibres. MT and F-actin recruitment is essential for FA organization and dynamics [20]. While FA assembly is actin-driven, disassembly requires their interaction with MTs and subsequent internalisation, resulting in contact dissociation from the ECM. Although Vimentin IFs have also been shown to directly associate with FA molecule Vinculin and control its localization, MT targeting of FAs is essential for FA disassembly and thereby cell migration. Gross disorganization of cytoskeleton had suggested defective adhesion of the KD cells to the ECM (Figure 1A, B). Expectedly, the bent and unaligned MTs in KD cells were unable to reach FAs as compared to the controls, which efficiently targeted FAs radially (Figure 4A). Immunolocalization of FA molecules Vinculin and Paxillin revealed a dramatic increase in size and reduction in number of FAs upon Rudhira depletion as compared to control, suggesting impaired FA dynamics (Figure 4B, S2A). This was confirmed by staining the cells with a phospho-tyrosine (pY) antibody as FA proteins are highly tyrosine-phosphorylated (Figure S2B). Further, immunoblotting showed that the levels of Vinculin and Paxillin were not significantly altered (Figure 4C). Conversely, Rudhira overexpression increases migration rate [11], and as expected, immunolocalization analysis showed a mild decrease in FA size in Rudhira overexpressing cells (Rudh2AGFP) as compared to the untransfected or vector controls (Figure S2C). Further, transient overexpression of Rudhira in KD cells rescued the FA size phenotype (Figure 4D). Therefore, we hypothesized that Rudhira depletion may increase FA assembly, or decrease disassembly or both. Double-immuno-localization showed that Rudhira does not co-localize with Paxillin or pY (Figure S2D, D’), suggesting that Rudhira controls cytoskeletal organization and dynamics resulting in modulated downstream FA dynamics and cell migration.

**Figure 4.**
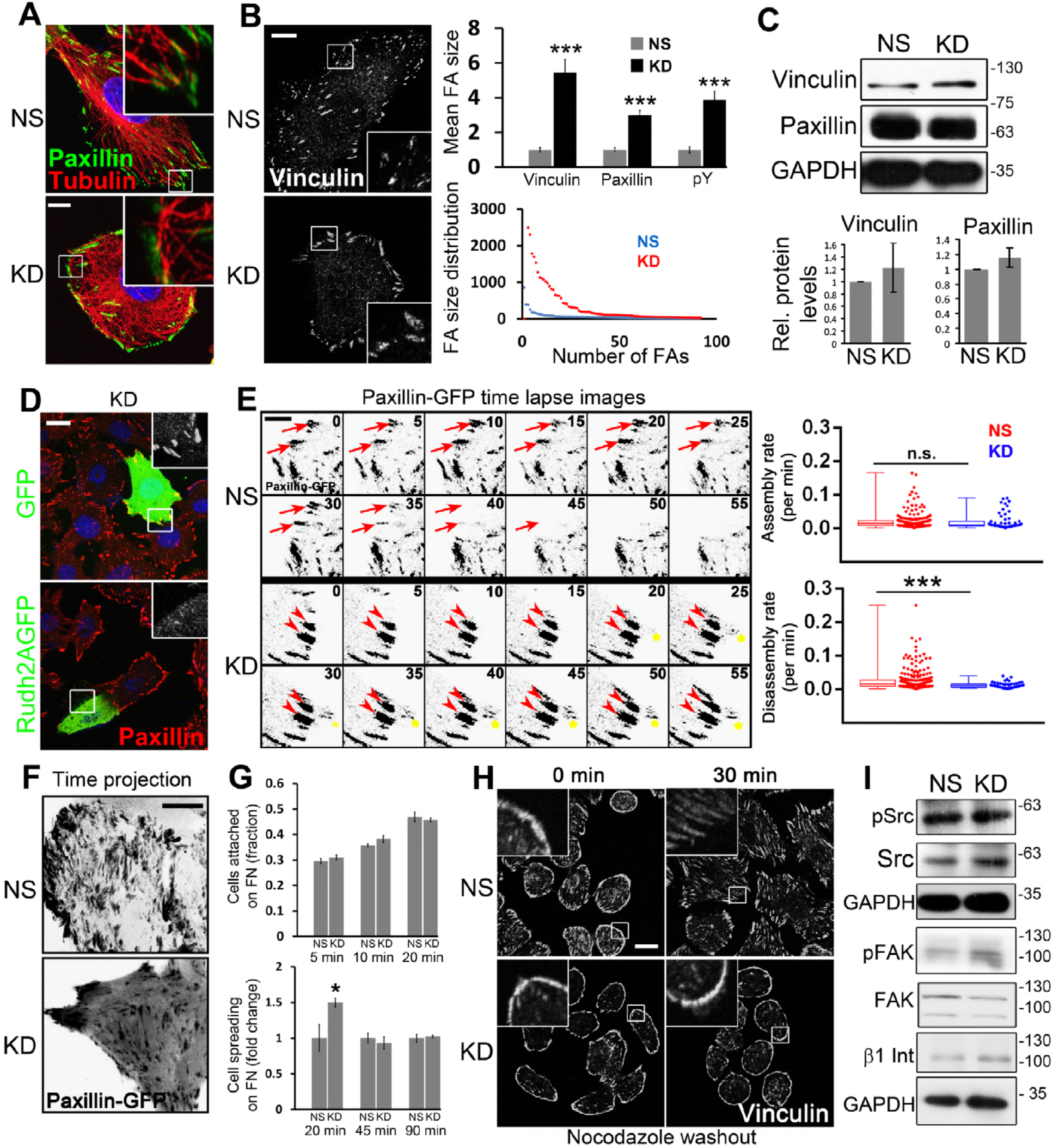
MT-mediated FA disassembly is impaired upon Rudhira depletion. (A) Immunostaining for Tubulin and Paxillin to detect the association of cell peripheral MTs with FA. Non-silencing control (NS) or *rudhira* knockdown (KD) endothelial cells (SVEC) were analyzed by immunostaining (B) or by immunoblot of cell lysates (C) to detect FAs marked by Vinculin, Paxillin and p-Tyrosine (pY) as indicated. Graphs show the quantitation of FA size, and the size and number distribution for FAs from 10, 8 and 6 images for analysis of Vinculin, Paxillin and pY respectively. (D) KD cells were transiently transfected with Rudhira-2A-GFP (Rudh2AGFP) or EGFP vectors and analyzed for FA size by immunostaining for Paxillin. (E) Time-lapse images of NS and KD cells transiently transfected with Paxillin-GFP monitored for 2-3 h and shown at 5-min intervals. Red arrow indicates a FA getting disassembled. Red arrowhead indicates a FA persisting over time and not getting disassembled. Yellow asterisk indicates a FA getting assembled over time. Graphs show the quantitation of FA assembly and disassembly rates computed using Focal Adhesion Analysis Server (FAAS) (see Methods), represented as whisker plots combined with scatter plots to show the distribution of individual FA. 5-7 optical slices were taken at 5-min intervals for 2 to 3 h. FAAS identified 225 and 240 FA in NS cells for assembly and disassembly respectively; and 57 and 58 FA in KD cells for assembly and disassembly respectively. 8-10 live cells each for NS and KD were imaged and analyzed. (F) Time-projected images of NS and KD cells imaged live after transient transfection with Paxillin-GFP and projected over 2 h to show the dynamics of FAs. (G) Quantitation of attachment and spreading profiles of cells on fibronectin with time, as indicated. (H) Recovery after Nocodazole treatment and immunostaining of fixed cells to detect FAs (marked by Vinculin). Cells were co-stained with Phalloidin (also see Figure S2A, B) to detect F-actin and DAPI to mark nuclei (Blue). Boxed regions in (A, B, D, H) are magnified in the insets. (I) Immunoblot of NS or KD cell lysates to detect the levels of FA signaling proteins Src, pSrc, FAK, pFAK and β1 Integrin. Error bars indicate standard error of mean (SEM). Results shown are a representative of at least three independent experiments. Statistical analysis was carried out using one-way ANOVA. Scale Bar: (A, B, F) 10 μm, (D, H) 20 μm, (E) 1 μm. *p<0.05, **p<0.01, ***p<0.001.

### Rudhira depletion impairs MT-dependent FA disassembly

Directional cell migration requires continuous coordinated removal and formation (turnover) of FAs at the leading edge and release of attachment at the rear. Defects in the process of FA assembly or disassembly are both detrimental to cell migration. We examined the steady state dynamics of FAs in control and KD cells transiently transfected with Paxillin-GFP using time lapse live imaging (Figure 4E, F and Video S5). Our observations and analysis of the time-lapse images by the FAAS (Focal Adhesion Analysis Server, see Methods) [21] showed that FA assembly was not affected upon Rudhira depletion, while disassembly was reduced to half of that in the control cells (Figure 4E and Video S5). Time-projection of the live images also showed highly dynamic FAs in control cells, while KD FAs appeared to be immobile (Figure 4F and Video S5).

To confirm these results, we used specialized molecular and cellular functional assays. Rudhira-depleted cells did not show significant difference in attachment but spread earlier than controls on fibronectin matrix, indicating that FA assembly is not impaired (Figure 4G). The early initial spreading of *rudhira* knockdown cells could be due to the persistence of FAs even after the 20 min in suspension, within which time FAs disassemble in control cells. Treatment with the MT depolymerising agent, Nocodazole inhibits FA disassembly as MTs are not recruited to FA [22]. Upon Nocodazole treatment, while control cells showed FA disassembly after 30 min of Nocodazole washout, Rudhira KD ECs continued to show large FAs that failed to turnover (Figure 4H and Figure S3A, B). Paxillin turnover is indicative of FA turnover. Upon Cycloheximide treatment, Paxillin levels dropped in control cells within 7 h (the half-life of Paxillin) but showed only minimal reduction in KD cells, suggesting impaired FA turnover (Figure S3C). Taken together these data suggest that Rudhira functions in MT-mediated FA disassembly. It is unlikely however, that Rudhira is a FA relaxing molecule, since MTs are dispensable for gross localization of Rudhira [11].

FA disassembly requires FA kinase (FAK) phosphorylation, subsequent MT-targeting of FAs and Dynamin2-mediated FA internalisation [22]. Immunolocalization and immunoblotting showed that Rudhira depletion does not alter levels of the components involved in FA-mediated signaling, namely FAK, pFAK and β1 Integrin, Src and pSrc and the early events of FA disassembly (Figure 4I, S3D, E). These data validate that Rudhira has a primary function at the cytoskeleton, downstream to which it promotes FA turnover and cell migration.

### Rudhira-dependent MT stability is essential for cytoskeletal organization

We next asked whether cytoskeletal crosstalk regulating MT stability was the primary mode of action of Rudhira. The Rho GTPase RhoA acts through Rho-associated Kinase (ROCK) to disassemble MTs and IFs. Treatment with ROCK inhibitor can restore MT assembly and stability, IF extension as well as FA dynamics. Rudhira KD cells treated with ROCKi showed almost complete rescue of phenotypes as observed by recovery of MT organization and cell-peripheral alignment and reduced FA size (Figure 5A). Further, cortical actin bundles and stress fibres were dramatically reduced (Figure 5A’). MTs did not bend and could reach the cell periphery, possibly because they were not impeded by the thick cortical actin (Figure 5A inset, A’). These data suggest that the primary function of Rudhira is to provide physiological stability to MTs, and the loss of Rudhira can be compensated for by stabilizing MTs or inhibiting MT disassembly pharmacologically. However, MTs may also reorganize in response to ROCKi-induced cell shape changes, which cannot be ruled out. It is also possible that Rudhira depletion deregulates Rho GTPase effectors like mDia and Tau, to affect MT stability.

**Figure 5.**
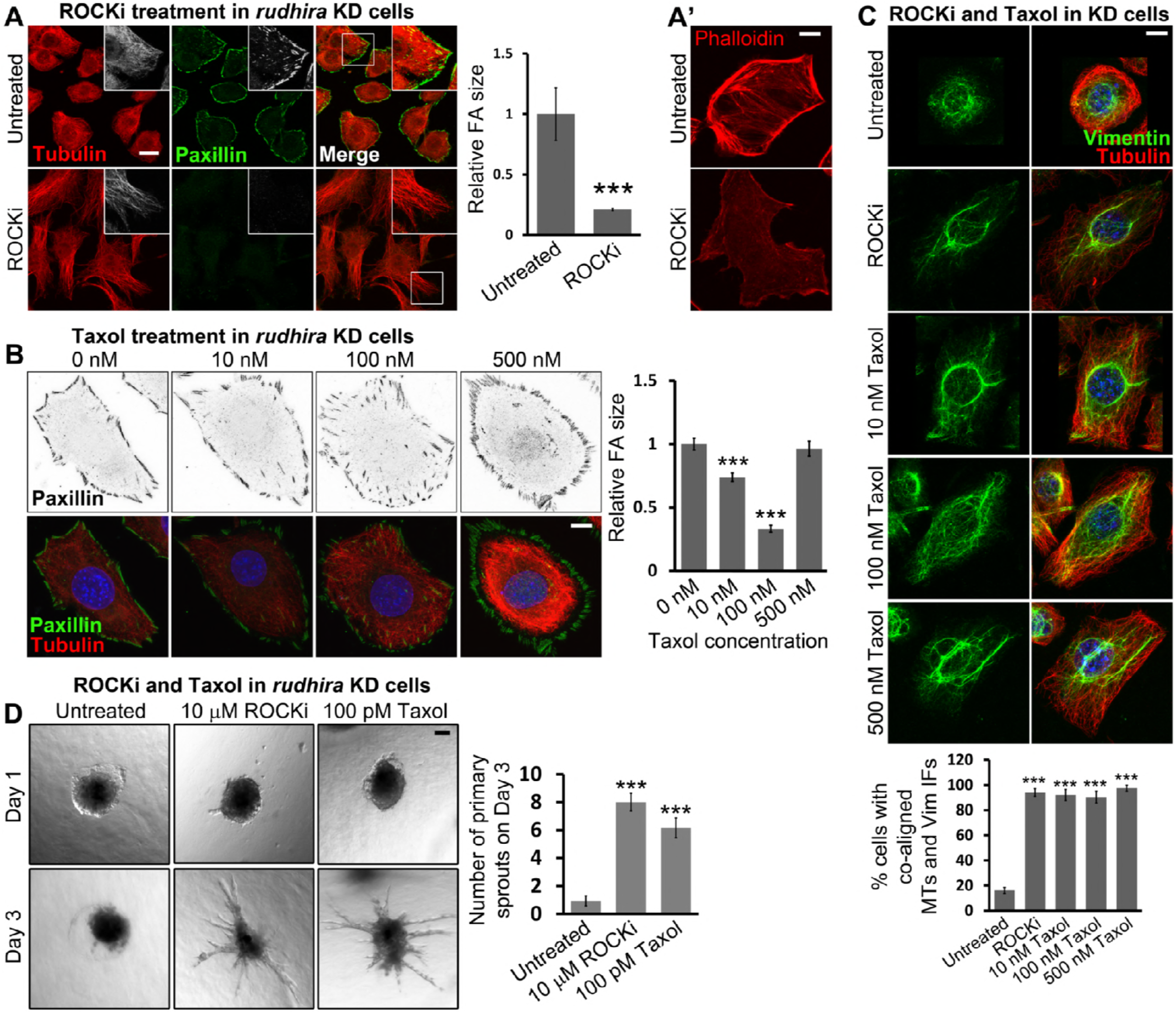
Rudhira stabilises microtubules for cytoskeletal organization and angiogenesis. (A, A’) Rudhira KD cells were kept untreated or treated with 10 μM ROCKi and analyzed for FA size and MT organization by co-immunostaining for Paxillin and Tubulin (A) or actin using Phalloidin (A’). Graph shows the quantification of relative FA size in ROCKi treated or untreated KD cells. Boxed regions in (A) are magnified in the insets. (B) Rudhira KD cells were treated with different concentrations of Taxol (Paclitaxel) as indicated, and analyzed for FA size and MT stabilization by co-immunostaining for Paxillin and Tubulin. Note the increasing fluorescence intensity of Tubulin and bundling (stability) of MTs with the increase in Taxol concentration. Graph shows the quantification of relative FA size in KD cells treated with different Taxol concentrations, compared to the untreated. (C) KD cells were treated with ROCKi or a range of Taxol concentrations, as indicated and analysed for MT-IF association by co-immunostaining for Vimentin and Tubulin. Graph shows the percentage of cells with coaligned MTs and IFs in each condition. (D) Spheroids formed from Rudhira KD cells were taken for collagen-based spheroid sprouting assay, in the presence or absence of ROCKi or Taxol, as indicated. Graph shows the quantification of the number of primary sprouts formed on Day 3 upon each treatment. Error bars indicate standard error of mean (SEM). Results shown are a representative of at least three independent experiments. Statistical analysis was carried out using one-way ANOVA. Scale Bar: (A) 20 μm, (A’, B, C) 10 μm, (D) 100 μm. *p<0.05, **p<0.01, ***p<0.001.

To dissect the effect of Rudhira depletion on MT stability from other properties leading to defective FA turnover, like cell shape changes, we transiently stabilized MTs in KD cells with Paclitaxel and scored for FA size. We observed a dose-dependent decrease in FA size with increasing concentration of Paclitaxel (Taxol, 10 nM to 100 nM) (Figure 5B). This suggests that drug-mediated MT stabilization can partially rescue the FA phenotype resulting from loss of Rudhira. However, at higher concentrations of Paclitaxel, FA size increased again, possibly due to drastic loss of MT dynamics, which could impede FA turnover. More interestingly, transient treatment with either ROCKi or Taxol led to the reorganization of Vimentin IFs and their co-association with MTs in KD cells (Figure 5C). Importantly, all concentrations of Taxol (10 nM to 500 nM) resulted in the extension of Vimentin IFs and their association with MTs. These data indicate that pharmacologically stabilizing MTs while still maintaining their dynamics in Rudhira depleted cells is sufficient to restore normal cytoskeletal organization. These data suggest that the primary role of Rudhira is to stabilize MTs *in vivo*, likely by crosslinking MTs and IF components in endothelial cells.

### Rudhira regulates MT stability for angiogenic sprouting

Rudhira functions in endothelial cell migration during angiogenesis. In mouse, loss of Rudhira causes mid-gestation lethality due to severe cardiovascular patterning defects and the loss of angiogenic sprouting [12]. As both MTs and Vimentin IFs have critical roles in endothelial sprouting [3], we hypothesized that the loss of sprouting in KD cells was due to a loss in MT-IF association and reduced physiological stability of MTs. Rho kinase inhibitor (ROCKi) and Taxol treatment are widely used to modulate sprouting angiogenesis [2, 23]. While Rho kinase (ROCK) inhibition promotes sprouting, low-dose Taxol (100 pM) is either inhibitory (in normoxia) or ineffective (hypoxia) in normal cells. Taxol treatment could stabilize MTs and promote MT-IF association at both high and low concentrations, like ROCKi treatment. However, for functional rescue we used low Taxol concentration as both the dynamics and the stability of MTs are essential for sprouting. KD cells, which otherwise fail to sprout as reported earlier [12], when treated with either ROCKi or low-dose (100 pM) Taxol rescued sprouting angiogenesis (Figure 5D). This suggests that Rudhira is essential for MT stability, cytoskeletal crosslinking, organization and dynamics during developmental vascular remodeling.

### The C-terminal BCAS3 domain is necessary and sufficient for cytoskeletal organization and cell migration

To elucidate how the organization of Rudhira protein mediates its function, we undertook a deletion analysis. Rudhira is reported to have predicted WD40-like structural domains, involved in protein interactions, at the N-terminal region and an uncharacterized BCAS3 domain in the C-terminal region [24]. Using multiple bioinformatics domain analysis servers and based on high confidence score we mapped the limits of these domains (Figure S4A, also see Methods). Rudhira also encodes multiple isoforms, and a shorter isoform of unknown function that lacks the initial 229 residues is reported. Protein structure prediction tool Phyre2 predicted one β-propeller (maximum 99.8% confidence and 17% identity) near the N-terminus (residues 92-434) (Figure S4B) and RaptorX predicted the presence of two β-propellers (residues 57-350, 351-582) (Figure S4B’). Interestingly, the C-terminal region did not align to any structure and is considered to be highly disordered (Figure S4B, B’). A PEST motif (signal for protein degradation, residues 883-903) was also identified in the C-terminal region (Figure 6A, S4A).

**Figure 6.**
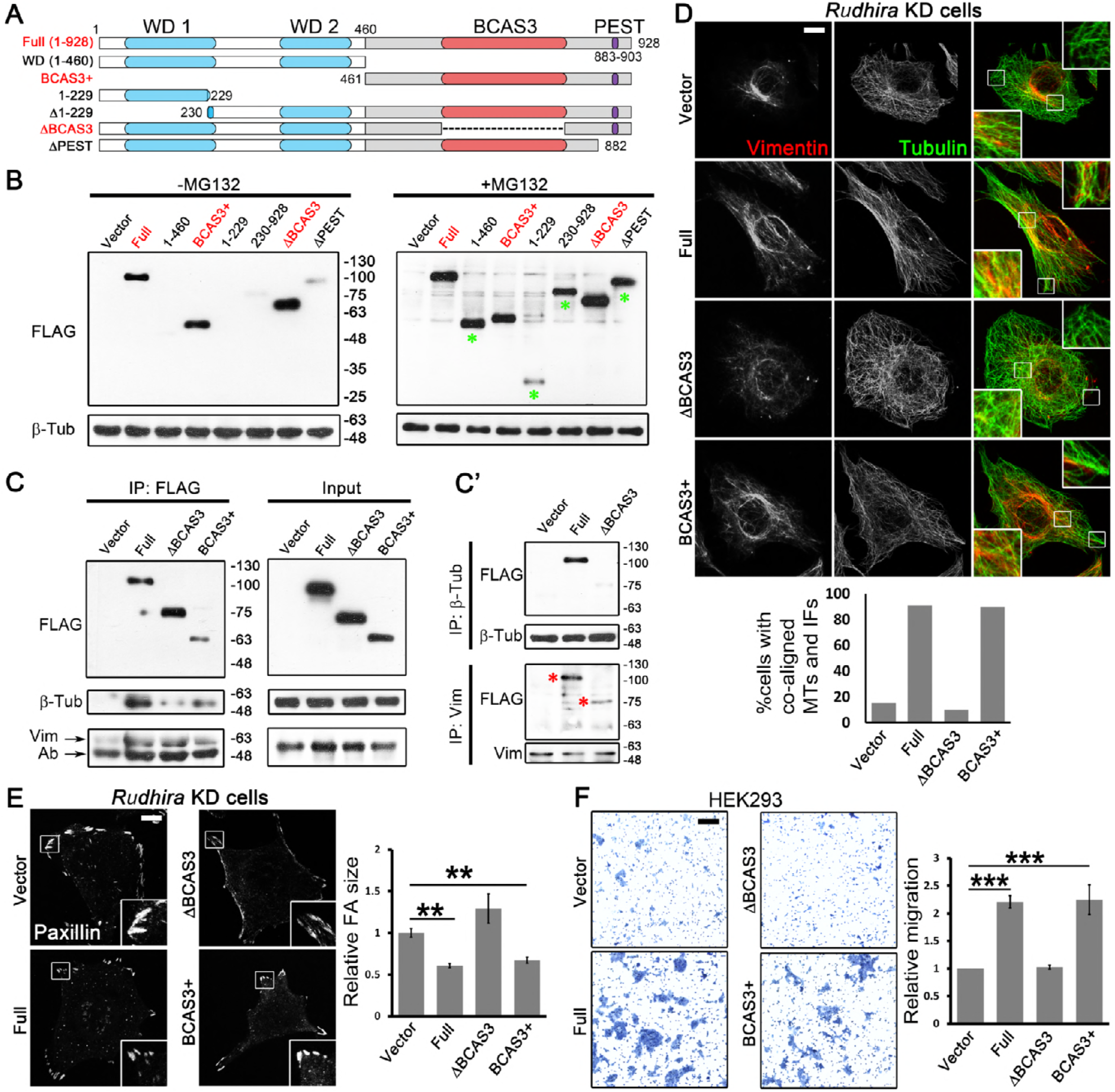
BCAS3 domain of Rudhira is necessary and sufficient for cytoskeletal crosstalk, organization and cell migration. (A) Schematic showing the deletion mutants of different regions of Rudhira protein, based on putative motifs/domains identified using bioinformatics analyses (see Methods). WD1, WD2: WD40 domains (blue); BCAS3: BCAS3 domain (orange); PEST: PEST motif (purple). Dotted line indicates region deleted in ΔBCAS3. (B) Validation of the expression of FLAG-tagged Rudhira full-length or deletion mutants overexpressed in HEK293T cells by immunoblot, with or without MG132, as indicated. (C, C’) FLAG-tagged Rudhira full-length BCAS3 or BCAS3+ or ΔBCAS3 fragments were analysed for interaction with β-Tubulin or Vimentin by co-immunoprecipitation with FLAG antibody, confirmed by reverse co-immunoprecipitation with β-Tubulin or Vimentin antibody (C’). (D, E) Rudhira full-length or fragments (cloned in pIRES2-EGFP vector) were transiently transfected in KD cell line to test for the rescue of MT, Vimentin IF organization, MT-Vimentin IF association and FA organization by double immunostaining for Tubulin and Vimentin (D), Tubulin and actin (see Supplementary Figure S4F) or FA marker Paxillin (E). Boxed regions in (D, E) are magnified in the insets and show perinuclear and cell peripheral regions (D) or only the peripheral regions (E). Graphs show the quantitation of percentage of cells showing coaligned MTs and Vimentin IFs from 10 cells and Paxillin FA size from at least 13 cells. (F) Rudhira full-length or fragments were overexpressed in HEK293 cells and tested for function using a transwell-migration assay, quantified in the graph. Error bars indicate standard error of mean (SEM). Results shown are a representative of at least three independent experiments. Statistical analysis was carried out using one-way ANOVA. Scale Bar: (D, E) 10 μm, (F) 100 μm. *p<0.05, **p<0.01, ***p<0.001.

While the WD40 domains function in protein-protein interactions, the BCAS3 domain is reported in proteins expressed in breast cancer and implicated in the progression of breast cancer [25]. Interestingly, a majority of the Rudhira post-translational modifications (PTMs) identified in high-throughput mass spectrometric screens were present in the C-terminal half of the protein (BCAS3+ fragment), which houses the BCAS3 domain, typically 229-245 amino acids long (Figure S4C). Analysis with MAPanalyzer (Microtubule-associated Protein Analyzer; http://systbio.cau.edu.cn/mappred/) [26] showed putative MT-interacting motifs in Rudhira distributed along the entire length of the protein (Figure S4D). Structural prediction using RaptorX suggested that while the N-terminal 1-460 fragment containing WD40 domains would form a 6-bladed β-propeller and ΔBCAS3 (Δ522-805) would form two β-propellers as in the full-length protein, the BCAS3+ fragment (461-928) would be mostly disordered (Figure S4E). Based on this information, we generated deletion mutants harbouring/lacking the putative domains or isoforms (Figure 6A) (see Methods) and expressed them in HEK293T cells.

Some of the deletion mutants expressed poorly, suggesting that the fragments may be unstable. Treatment with the proteasomal inhibitor MG132 stabilized these fragments (Figure 6B). However, the WD40 domain containing fragment devoid of the BCAS3 domain (ΔBCAS3), and the BCAS3 domain containing fragment (BCAS3+), both expressed at levels similar to those of the full-length protein (Full). Also, the WD40 and the BCAS3 domains were predicted at high confidence by multiple bioinformatics tools, as compared to other domains/motifs (Figure S4A). Hence further molecular and functional analysis was limited to these to avoid the possible differences due to varied expression levels.

Interestingly, the full-length protein as well as the BCAS3+ fragment could efficiently co-immunoprecipitate tubulin. However, tubulin interaction was dramatically reduced with ΔBCAS3 (Figure 6C), suggesting that the BCAS3 domain of Rudhira and not the WD40 domain, is necessary and sufficient for Tubulin/MT-interaction. In addition, all the three fragments could co-immunoprecipitate Vimentin. This was confirmed by reverse co-immunoprecipitation with Tubulin and Vimentin, wherein Tubulin could efficiently pull-down the full-length protein but not ΔBCAS3 mutant while Vimentin could pull-down the full-length as well as ΔBCAS3, although the interaction with ΔBCAS3 was slightly reduced as compared to the full-length (Figure 6C’). This suggested that while Rudhira contained multiple Vimentin-binding regions, Tubulin-binding regions were present mainly in the BCAS3+ fragment. It is important to note that, in ΔBCAS3, the region from 527 to 805 was deleted (instead of 521-792), because of the high sequence conservation in the BCAS3 domain till the 805 residue and to avoid the deletion of the overlapping SxIP motif (518-521). Also, the BCAS3+ fragment (461-928) had WD40 domains deleted but retained the BCAS3 domain and phosphorylation sites reported at high frequency.

To test whether the interactions with Tubulin and vimentin correlated with function, we overexpressed Rudhira full length or deletion mutants in KD cells and assayed for rescue of the KD phenotypes, namely reduced MT-Vimentin IF association, enlarged FAs and increased actin stress fibres. The full-length protein or BCAS3+ restored MT and Vimentin organization, MT-Vimentin IF association (Figure 6D), FA size and actin organization (Figure 6E, S4F, G, Video S6). However, ΔBCAS3-expressing KD cells continued to show disorganized actin and MTs, reduced and less extended Vimentin IFs (Figure 6D), loss of IF-MT alignment and large FAs (Figure 6E, S4F, G, Video S6). To test the functional relevance of the BCAS3 domain of Rudhira we checked the effect of its presence on cell migration in a trans-well assay. Overexpression of the full-length protein or the BCAS3+ in HEK293 cells resulted in an increase in migration, while ΔBCAS3 did not (Figure 6F). Together, these data show that Rudhira-cytoskeleton interactions leading to MT-IF crosstalk mediated by the BCAS3 domain is essential for regulating cytoskeleton architecture. Further, the BCAS3 domain is necessary and sufficient for Rudhira function.

## Discussion

In a dynamically regulated system such as the vasculature, controlled endothelial cell migration and sprouting angiogenesis are key to ensuring blood supply during development and tissue repair. Defects in this process can lead to developmental anomalies or even embryonic death. The cycle of endothelial cell proliferation, migration and sprouting angiogenesis is influenced by several physiological and pathological cues. Cytoskeletal remodeling underlies all of these processes and mediates molecular crosstalk to ensure a calibrated response over a range of signals. Cell type-specific components ensure an appropriate response to the dynamic cues from circulation as well as the tissue microenvironment. Rudhira is a dynamically regulated molecule with tissue-specific roles in regulating the cytoskeleton in endothelial migration and sprouting angiogenesis [11, 12]. Here we investigated the molecular mechanism by which Rudhira regulates the cytoskeleton and found that Rudhira crosslinks IF and MT cytoskeleton, stabilizes MTs and directs MTs for FA disassembly, mediated by its BCAS3 domain.

Vimentin IFs template MT growth and stabilize MTs. The presence of Rudhira, MTs and Vimentin IFs together suggests a major role of Rudhira in cytoskeletal crosstalk. The primary function of Rudhira appears to be binding to MTs and Vimentin IFs, providing physiological stability to MTs and promoting Vimentin IF extension, to aid cytoskeletal organization and downstream processes. It is, however possible that the restoration of cytoskeletal architecture and sprouting observed upon treatment with ROCKi or Taxol was not through their effect on MTs but rather due to probable effects on other cytoskeletal components including Rudhira, Vimentin or actin. We reported earlier that *rudhira* KD cells have dramatically reduced soluble Vimentin [12]. ROCKi and Taxol are also known to solubilize Vimentin, which may also rescue the loss of Rudhira. It is also possible that ROCKi or Taxol treatment may stabilize MTs or prevent their disassembly and simultaneously lead to the formation of secondary sites for MT nucleation, which may lead to the rescue of Rudhira depletion phenotypes and sprouting. Cytoskeletal crosstalk is complex, and it is likely and probable that Rudhira functions with other cytolinkers and cytoskeleton-associated molecules for coupling MTs and IFs.

Rudhira is developmentally essential and transiently expressed in angiogenic endothelium during vascular development. It will be interesting test the possibility of transient expression of Rudhira or a Rudhira-like molecule in other tissues which undergo dynamic remodeling. The cytolinker function of Rudhira sufficiently explains the molecular and cellular phenotypes observed upon its depletion. Identification and detailed characterization of the loss of function mutants of Rudhira or other cytolinkers may further our understanding and targeting of cytoskeletal crosstalk in vascular development and disease. Loss of Rudhira results in embryonic lethality in mouse with gross cardiovascular patterning defects. It is unlikely that stabilizing MTs would completely override the effect of loss of Rudhira. However, controlled restoration of MT stability and dynamics or the expression of another cytolinker in *rudhira* knockout may be useful in delineating the primary molecular function of Rudhira *in vivo*.

Aberrant cell-matrix adhesion is the underlying cause of defective migration in a variety of contexts [27]. FA turnover is a complex process. Molecules that regulate FA components are known, however local interactions that direct MTs to FAs remain unclear. +TIP proteins such as CLASPs localize to FAs as well as MTs, thereby bridging the two for FA disassembly [28]. Rudhira, on the other hand, binds MTs and IFs but not FAs, highlighting its role as a targeted regulator of the cytoskeleton in this process. The effect of Rudhira depletion on FA disassembly and thereby cell migration hence appears to be downstream of its more direct role in aligning growing MTs towards the cell periphery and mediating their coalignment with IFs thereby stabilizing them. Hence the role of Rudhira is to organize the MT cytoskeleton downstream of FAK phosphorylation to bring about FA disassembly. In addition, the unaligned growth and fewer growing MTs in KD cells suggest a role for Rudhira before MTs encounter actin stress fibres at the cell periphery. Rudhira-mediated control of FA and MT dynamics are unlikely to be independent, owing to the strong correlation between the organization of the two as suggested in the literature as well as in our study [10].

MT-growth initiates primarily from the MT organizing centres (MTOCs). Loss of, or structural defects in the MTOCs, which are primarily centrosomes in endothelial cells, may also explain the defects in MT growth and stability. However, earlier studies suggest that although the MTOC does not realign along the direction of migration, it is indeed present in *rudhira* KD cells [11]. The contribution from the defects in polarity caused by Rudhira depletion may also lead to disorganized MT growth and architecture [11]. Additionally, the involvement of independent molecular pathways controlling FA and MT organization cannot be ignored. Identification of further molecular interactors of Rudhira will delineate its position in the molecular pathway governing MT growth and recruitment to FAs. Bioinformatics analysis reveals the presence of SxIP motifs towards the C terminus of Rudhira, suggesting an interaction with EB proteins [29]. Thus, Rudhira may have a prominent role in MT growth towards FAs via EB proteins, known to be essential for MT growth and polarity. This also raises the interesting possibility that Rudhira may lay down tracks for MT growth.

The Rudhira protein has several conserved domains such as WD40 domains and the BCAS3 domain, however the relevance of its organization in the normal in vivo function of Rudhira is not known. WD40 domain-containing proteins assume a beta propeller structure that is thought to act as a scaffold for multiple protein interactions. However, our experiments suggest that the C terminal fragment bearing the BCAS3 domain, rather than the N-terminal WD40 domain containing fragment, could bind tubulin and importantly, restore function. In addition to the BCAS3 domain, the BCAS3+ fragment includes a PEST domain, SxIP motifs, some of the C-terminal region and frequently reported phosphorylation sites, which may also contribute to the function. However, these additional features require the BCAS3 domain, as removing the BCAS3 alone caused loss of BCAS3+ function. Some of the fragments, including the alternative isoform, the PEST motif deletion and others, showed increased susceptibility to ubiquitin-proteasome mediated degradation, suggesting complex regulation. It is possible that some of these, like the alternative isoform, could be transiently and dynamically expressed for temporal control of Rudhira function, a possibility that merits further investigation. It is likely that this isoform has a physiologically relevant role, which enhances Rudhira function in specific contexts, as it contains the conserved BCAS3 domain but lacks the WD40 or other domains. This report also assigns molecular function to the conserved BCAS3 domain sequence. It will be interesting to test whether the BCAS3 domains present in many autophagy-related proteins and proteins expressed in cancers share similar functions.

The cytoskeleton is involved in multiple processes. The restricted expression of Rudhira may permit context-dependent regulation of these processes. Rudhira/BCAS3 is implicated in metastatic carcinomas [13, 25], where MTs undergo differential association with MT-associated proteins and transition to dynamic instability and Vimentin IFs are upregulated. Hence, we speculate that Rudhira may also play a role in mitosis. During development, Rudhira may control MT stability and alignment and MT-IF association, thereby maintaining cell and tissue polarity as well as migration. Our studies will aid in revealing the dynamics of the interaction between MTs, IFs and Rudhira in various contexts where intricate association of cytoskeleton, FA remodeling and cell migration are essential.

Rudhira expression in endothelial cells and its effects on angiogenesis shows its key role in vascular development by the control of MT stability and cytoskeleton organization. The mis-expression of Rudhira/BCAS3 in grade III glioblastomas and other cancers and association with coronary artery disease make it a principal target in these diseases. Our finding, that the BCAS3 domain is required for promoting cytoskeleton crosstalk, and maintaining MT architecture and FA dynamics, will help devise strategies for controlled alteration of cytoskeletal architecture to correct aberrant cell migration, tissue malignancy or degeneration.

## Materials and methods

### Cell culture

Mouse Saphenous Vein Endothelial Cell line (SVEC) was obtained from Kaustabh Rau, National Centre for Biological Sciences, Bangalore and HEK293, HEK293T cells were from ATCC. Cells were cultured in DMEM (ThermoFisher Scientific, USA) supplemented with 10% FBS (Gibco, ThermoFisher Scientific, USA). Generation of knockdown lines is reported elsewhere [12]. Serum starvation was performed for 48 h in DMEM.

### Immunostaining, antibodies and small molecule treatment

Cells were fixed in 4% paraformaldehyde at room temperature for 15 min or 100% methanol at -20 °C for 10 min and processed for immunostaining using standard procedure. Primary antibodies used were against Rudhira [11], Vinculin, α-Tubulin, Ac-Tubulin (Sigma Chemical Co., USA) β-Tubulin (Developmental Studies Hybridoma Bank (DSHB), Iowa; ThermoFisher Scientific, USA; Abcam, USA), Paxillin, FAK, β1 Integrin (Merck, USA), Vimentin, Glu-Tubulin, EB1 (Abcam, USA), Plectin (Santa Cruz Biotechnology, USA), GFP (ThermoFisher Scientific, USA), pFAK, pY (Cell Signaling Technologies, USA). Secondary antibodies were coupled to Alexa-Fluor 488 or Alexa-Fluor 568 or Alexa-Fluor 633 (Molecular Probes, USA). Phalloidin was conjugated to Alexa-Fluor 633 (Molecular Probes, USA). Nocodazole, Cycloheximide, ROCK inhibitor (ROCKi, Y27632) and Taxol (Paclitaxel) were from Sigma Chemical Co., USA. NS and KD cells were treated with 50 μg/ml of Cycloheximide for a period of 0, 7 and 14 h. Thereafter, the cells were taken for immunoblot analysis (Figure S3C). Cells were treated with ROCKi or Taxol for 1 h and processed for immunostaining with Paxillin, Tubulin or Vimentin antibodies or Phalloidin, as indicated (Figure 5A, B, C).

### Fluorescence microscopy, live cell imaging and analysis

Confocal microscopes (LSM 510 Meta, LSM 880 with Airy Scan from Zeiss, FV3000 from Olympus), Spinning Disc Microscope (Perkin Elmer with Yokogawa camera attachment) or a motorized inverted microscope with fluorescence attachment (IX81, Olympus) were used for fluorescence microscopy and time lapse imaging. Line profile of fluorescence intensities (Figure 3B, 4F) was generated in ZEN Blue software from Zeiss. Super-resolution microscopy was performed by imaging in the Airy Scan image acquisition and processing mode of the LSM 880, Zeiss. For live cell imaging, a sample heater (37°C) and CO_2_ incubation chamber (Tokai Hit) were used to control temperature and CO_2_ levels during live cell imaging. All images in a set were adjusted equally for brightness and contrast using Adobe Photoshop CS2, where required. Rudhira NS or KD cells were transiently transfected with EB1-GFP and seeded on fibronectin-coated glass-bottom dishes. Live imaging for EB1-GFP was carried out 24 h post seeding for 3 min at 4 s intervals to determine MT growth and alignment. EB1-GFP live images were time-projected in ImageJ (NIH) to represent MT growth. EB1-GFP tracks were generated manually and residence time was calculated manually using ImageJ (NIH) with the manual tracking plug-in. Rudhira NS or KD cells were transiently transfected with Paxillin-GFP and seeded on fibronectin-coated glass-bottom dishes. Live imaging for Paxillin-GFP was carried out 24 h post seeding for 2-3 h at 5 min intervals to determine FA assembly and disassembly rates under steady state. The images were processed for estimation of various parameters using Focal Adhesion Analysis Server (FAAS) [21]. Paxillin-GFP, EB1-GFP live images were time-projected in ImageJ (NIH) to represent FA and MT growth dynamics respectively. Rudhira NS or KD cells were transiently transfected with Vimentin-GFP and seeded on fibronectin-coated glass-bottom dishes. Cells were incubated with 250 nM SiR-Tubulin for 2 hours before live imaging was carried out for 4 min at 10 s intervals to test coalignment of MTs and IFs.

### *In situ* Proximity Ligation Assay (PLA or Duolink assay)

*In situ* PLA (Proximity Ligation Assay) reaction was performed on SVEC cell lines. The cells were cultured, fixed, permeabilised and stained with primary antibodies as indicated. Thereafter, the protocol for PLA as recommended by manufacturer (Duolink, USA) was followed. Post PLA, nuclei were counterstained with DAPI.

### Cell attachment and spreading assays

The assays and quantitation were carried out as mentioned in [30] which has cited [31], with a few modifications. Briefly, 96-well plates were coated with fibronectin (10 μg/ml) for 60 min at room temperature and blocked with heat-denatured filter-sterilized BSA for 30 min at room temperature. Cells were put in suspension in warm medium at 37 °C, 5% CO_2_ to disassemble already formed FA. Thereafter, 20000 (for attachment) or 10000 (for spreading) cells were seeded per well and allowed to attach or spread for the indicated times. Floating or loosely attached cells were removed by washing twice with PBS and then fixed with 4% paraformaldehyde. For spreading assay, the extent of spreading was quantified in ImageJ (NIH) from RFP (expressed from the NS or KD vector) images. For attachment assay, cells were stained with 0.1% crystal violet for 60 min at room temperature, washed three times with water and the dye was solubilised in 100 μl 10% acetic acid and the absorbance was measured at 570 nm using a plate reader. The total number of cells attached at 3 h was set to 100% for both NS and KD lines.

### Nocodazole and cold treatment

For MT stability experiments (Figure 3C, C’, D, S4B), cells were treated with indicated dosages of Nocodazole or cold Phosphate buffered Saline (PBS) (4 °C) for 30 min, washed twice with cold PBS and fixed in ice-cold methanol at -20 °C. For MT recovery experiments (Figure 2H, S3A, B) cells were treated with Nocodazole (10 μM) for 30 min in complete medium at 37 °C, 5% CO_2_, washed twice with PBS and incubated with fresh culture medium for desired time intervals as indicated. Thereafter, cells were fixed and taken for immunostaining of Vinculin, F-actin (Phalloidin) and α-Tubulin, as indicated.

### SiR-Tubulin labelling of MTs

Silico-Rhodamine conjugated Docetaxal (SiR-Tubulin) was as used in [32] and was a kind gift from Sarit Agasti, JNCASR. MT labelling was performed as indicated in [32]. Briefly, cells were incubated with 250 nM of SiR-Tubulin for 2 h in complete medium at 37 °C, 5% CO_2_, washed twice with fresh culture medium and taken for live cell imaging.

### Co-immunoprecipitation and Western blot analysis

50 μg lysate was used for Western blot analysis by standard protocols. Primary antibodies used were as indicated earlier. HRP-conjugated secondary antibodies against appropriate species were used and signal developed using Clarity Western ECL substrate (Biorad, USA). Western blot intensities were normalised to GAPDH and quantification was carried out using ImageJ (NIH). For co-immunoprecipitation assays, 500 μg lysate of HEK293T cells overexpressing Rudhira fragments was incubated overnight with 10 μl of FLAG M2 beads (Sigma Chemical Co., USA) or 10 μl of β-Tubulin antibody (DSHB, Iowa) or Vimentin antibody (Sigma Chemical Co., USA), captured on Protein G-sepharose beads (Sigma Chemical Co., USA), washed three times in lysis buffer and analyzed by immunoblotting with anti-β-tubulin (Abcam, USA), Vimentin (Sigma Chemical Co., USA; Abcam, USA) or FLAG (Sigma Chemical Co., USA) antibody.

### Spheroid sprouting and transwell migration assay

The assays and quantitation were carried out as described previously [11] [12]. Briefly, for spheroid sprouting, 750 cells each of the KD line were taken for spheroid formation in a round-bottom non-adherent 96-well dish (Costar, USA), in 1% CMC (carboxy methyl cellulose) in 10% FBS in DMEM. The spheroids formed were transferred to collagen gels (Rat tail, Type I, ThermoFisher Scientific, USA) with a final concentration of 2.5 mg/ml, with or without ROCKi or Taxol. Gels were overlaid with 200 μl of 10% FBS in DMEM and the sprouting was monitored for 3 days. For transwell migration, 24 h after transfection with desired plasmid vectors, cells were serum-starved for 12 h and 20000 cells were plated onto the upper chamber of the transwell filter inserts with 8 μm pore size, 24-well format (Costar, USA). 10% serum medium was added to the lower chamber to serve as a chemo-attractant. After 24 h, cells were fixed in 4% paraformaldehyde for 10 min at room temperature. Cells on the top of the filter were removed using a cotton swab. Cells that had migrated to the bottom were fixed and stained with 0.5% crystal violet for 10 min at room temperature. The dye was extracted in methanol and absorbance measured spectrophotometrically at 570 nm.

### Rudhira *in silico* analysis, deletion mutant cloning, plasmid constructs and transfection

Domain/motif prediction analysis of mouse Rudhira protein sequence (Uniprot Id Q8CCN5.2) was performed using various bioinformatics tools, namely Superfamily (http://supfam.org/SUPERFAMILY/), Motif Scan (https://myhits.isb-sib.ch/cgi-bin/motif_scan), Pfam (https://pfam.xfam.org/), NCBI-CDD (https://www.ncbi.nlm.nih.gov/Structure/cdd/wrpsb.cgi?INPUT_TYPE=precalc&SEQUENCE=33300978), Motif Finder (http://www.genome.jp/tools/motif/), Interpro (https://www.ebi.ac.uk/interpro/), epestfind in the EMBOSS package of ExPASy (https://www.expasy.org/tools/). Post Translational Modifications (PTMs) in Rudhira were identified using PhosphoSitePlus (https://www.phosphosite.org/homeAction.action). Microtubule binding regions were predicted using MAPanalyser (http://systbio.cau.edu.cn/mappred/). Rudhira full length protein or deletion mutant structure prediction was performed using Phyre2 (http://www.sbg.bio.ic.ac.uk/phyre2/html/page.cgi?id=index) and RaptorX (http://raptorx.uchicago.edu/).

pCMV-Rudh-IRES2-EGFP, pCAG-Rudh-2A-GFP (Figure 2D, S2C), pCMV-RudhFL-FLAG were described earlier [11]. Rudhira ORF was sub-cloned from pCAG-Rudh-2A-GFP vector into pEGFP-N3 vector (Clontech, USA) using NheI-SacII sites to obtain pCAG-Rudhira-GFP (Figure 3D, S4A, B). For deletion mutant cloning, the regions to be cloned were PCR amplified from pCMV-RudhFL-FLAG vector and cloned in pCMV-Tag2B vector (Stratagene, USA) (Figure 5B, C). The desired fragments from pCMV-Tag2B vectors were digested using EcoRI-XhoI and sub-cloned into the compatible EcoRI-SalI sites of pIRES2-EGFP vector (Clontech), to obtain GFP fluorescent reporter plasmids (Figure 3E, 5D, E, F, S5F, G). EB1-GFP plasmid was a kind gift from Yuko Mimori-Kiyosue (Riken Kobe, Japan) and Paxillin-GFP was a kind gift from Rick Horwitz. HEK293 and HEK293T cells were transfected using Calcium Phosphate method and SVEC cells were transfected using Lipofectamine 2000 (ThermoFisher Scientific, USA).

### Quantification and Statistical analyses

Statistical significance analyses were performed using One Way ANOVA in the Data Analysis package in Microsoft Excel. p<0.05 was considered significant.

## Acknowledgements

We thank Yuko Mimori-Kiyosue, Riken Kobe, Japan for EB1-GFP plasmid, Rick Horwitz for Paxillin-GFP plasmid, Sandrine Etienne-Manneville, Institut Pasteur, Paris for Vimentin-GFP plasmid, Sarit Agasti and Ranjan Sasmal, JNCASR for SiR-Tubulin; Preeti Jindal, Abarna Sinha and Aksah Sam for Bioinformatics analyses and generating deletion mutants; JNCASR Imaging facility, NCBS Central Imaging and Flow Facility, and laboratory members for fruitful discussions. This work was funded by grants from the Department of Biotechnology, Government of India (Sanction no. BT/PR11246/BRB/10/644/2008 dated 29.09.2009) the Wellcome Trust, UK (094879/B/10/Z) and intramural funds from JNCASR, India. Maneesha S. Inamdar conceived of the project and directed the work. Maneesha S. Inamdar, Divyesh Joshi designed and performed all experiments, wrote the manuscript. The authors declare that they have no conflict of interest. Student’s t-test or One Way ANOVA was used for statistical significance. All relevant data are within the paper and its Supplementary Information files.

## Abbreviations

Rudh: Rudhira
BCAS3: Breast Carcinoma Amplified Sequence 3
MT: Microtubule
IF: Intermediate Filament
Tub: Tubulin
Vim: Vimentin
FA: Focal Adhesion
GFP: Green Fluorescent Protein
EB1: End Binding Protein 1
Minute: min
Hour: h
Second: s

